# Pioneer factor FOXA1 can boost CHO cell productivity

**DOI:** 10.1101/2025.09.10.675314

**Authors:** Sienna P. Butterfield, Fay L. Saunders, Robert J. White

## Abstract

Industrial production of biologics commonly involves the integration of transgenes into genomes of host cells, such as Chinese hamster ovary (CHO) cells. A major determinant of productivity is the epigenetic control of the transgene promoter, accessibility of which can decrease during production due to the spread of heterochromatin. Pioneer factors such as forkhead box A1 (FOXA1) can bind heterochromatin, increase its accessibility and facilitate transcription of target genes. We show that FOXA1 can bind the EF1α promoter, which is widely used in industry. Overexpressing FOXA1 in an industrially-relevant CHO-DG44 cell line raised production of monoclonal antibody encoded by transgenes transcribed from EF1α promoters. Mechanistically, this response can be attributed to recruitment by FOXA1 of epigenetic modifiers and a chromatin remodelling complex, which reprogramme the promoter to optimise transcription. In parallel, FOXA1 overexpression induces endogenous genes with beneficial effects on cell viability. This strategy significantly enhanced cell-specific productivity, demonstrating potential benefit in biomanufacturing.

## 1 Introduction

Extensive optimisation of CHO cell lines and growth conditions has enabled industrial mAb production titres to exceed 10 g/L.^[1,2]^ However, unpredictable epigenetic changes over production timelines remain a persistent challenge, causing expression levels to fall in initially high-producing clones.^[3]^ Epigenetic modifications, including hypermethylation of promoter DNA and deacetylation of histones, can incorporate transgenes into compacted heterochromatin that is less accessible to transcription machinery.^[4,5]^ Current methods to combat such silencing include the use of barrier elements to protect transgenes and maintain expression stability.^[6,7]^ Targeted epigenetic editing is an alternative, which can be achieved using CRISPR/dCas systems to deliver synthetic derivatives of epigenetic modifiers, such as histone acetyltransferases (HATs).^[8–11]^ However, expression of these large synthetic proteins increases the transcriptional burden of production cell lines.^[10]^

Pioneer transcription factors can bind and open condensed chromatin.^[12]^ FOXA1 was the first example to be characterised and was shown *in vitro* to bind DNA wrapped around nucleosomes.^[13]^ Footprinting *in vivo* revealed that FOXA1 occupies its target loci prior to their expression, establishing a model in which FOXA1 binds closed, inactive chromatin to induce changes that allow transcription.^[12,14]^ The DNA-binding domain of FOXA1 contains three α-helices, three β-strands and two peripheral wing loops.^[15–17]^ Nucleosomal binding is driven predominantly by non-specific interactions made via the wing domains, whilst the α-helices bind along the major groove of DNA, making base-specific contacts.^[18]^

FOXA1 can displace linker histone H1,^[19,20]^ thereby reducing chromatin compaction and improving accessibility.^[20,21]^ It can then recruit remodelling complexes, such as SWI/SNF, which can open chromatin by moving nucleosomes.^[22,23]^ FOXA1 also contributes to epigenetic signatures associated with active transcription, including DNA hypomethylation and histone H3 lysine 27 (H3K27) acetylation.^[24]^ For example, FOXA1 forms a feed-forward regulatory loop with the 5-methylcytosine dioxygenase TET1, both inducing its expression and recruiting it to target sites.^[25]^ Gene activation by FOXA1 involves demethylation of DNA by TET1.^[24,26]^ FOXA1 can also bind and recruit histone acetyltransferase p300, leading to gene activation through H3K27 acetylation.^[27]^ By thus reprogramming the epigenetic landscape, FOXA1 can stimulate transcription of target genes.^[28,29]^

FOXA1 overexpression was found to stimulate mAb production in CHO-M cells, a CHO-K1-derived cell line adapted for suspension culture.^[30]^ This effect did not involve induction of transgene mRNAs, but was instead achieved indirectly via stimulation of endogenous host cell genes, including the Tagap and Ca3 genes that are involved in cytoskeleton remodelling and resilience to stress, respectively.^[30]^ Indeed, overexpression of Tagap or Ca3 was sufficient to stimulate mAb production in CHO-M cells.^[30]^

We present evidence that FOXA1 can also stimulate the endogenous Tagap and Ca3 genes when overexpressed in the Apollo^TM^ platform, a CHO-DG44-derived expression system that is extensively used in industry. We show that FOXA1 binds directly to these genes and causes promoter demethylation. In addition, we demonstrate direct binding of FOXA1 to the EF1α promoter, which we use to drive transcription of mAb transgenes. Several epigenetic changes are identified at the transgene promoters, as well as the robust induction of mRNA output. These combined effects result in highly significant elevation of titre and specific productivity in a CHO cell platform that is optimised for industrial use.

## 2 Materials and Methods

### 2.1 Bioinformatic analysis

#### 2.1.1 Motif enrichment analysis

Motif enrichment analysis investigations were conducted using Ciiider (available at https://ciiider.com/), to identify transcription factor binding sites. Ciiider was supplied with the human EF1α promoter, CHO Tagap (NCBI 100761957) and CHO Ca3 (NCBI 100774549) sequences.^[31]^ FOXA1 motifs were obtained via JASPAR (available at https://jaspar.elixir.no/) in MEME format.^[32]^ Ciiider identified FOXA1 binding motifs within the provided sequence.

### 2.2 Cell culture

#### 2.2.1 Apollo cells

Clonal cell lines of recombinant Apollo cells, an industrially-relevant CHO-DG44 expression system (FUJIFILM Biotechnologies), were cultivated in CD OptiCHO^TM^ medium (Thermo Fisher) with 8 mM glutamine (Sigma Aldrich) at 37 °C, 8.0 % CO_2_. Cells stably expressed a construct containing Herceptin light and heavy chain genes driven by identical EF1α promoters. Cells were maintained by seeding every 2-3 days at 0.5×10^6^ cells/ml and counted using the Vi-CELL XR Cell Viability Analyser (Beckman Coulter).

#### 2.2.2 Overexpression of FOXA1

Apollo cells were transfected with either a plasmid encoding human FOXA1 linked to an HA-tag (FOXA1, Sino Biological Inc.) or a corresponding control plasmid (control, Sino Biological Inc.). For transfection, 0.5×10^6^ cells were pelleted and resuspended in 100 μl transfection buffer (Lonza) and transferred to a sterile 3 mm Amaxa nucleofection cuvette. Plasmid DNA (8 µg) was added dropwise, and cells were nucleofected using the U-26 programme on the Nucleofector IV (Lonza). Sterile culture medium (500 μl) was added and cells left for 15 mins before transferring to a 15 ml flask and left static at 37 °C, 8.0 % CO_2_. After 48 hrs, flasks were placed under agitation and for stable selection, media was supplemented with puromycin at 250 µg/ml every 2-3 days for 3 weeks.

### 2.3 Western blot analysis

2×10^6^ cells were isolated from culture supernatant, lysed in Laemmli SDS sample buffer (Bio-Rad) and β-mercaptoethanol (9:1 ratio) and boiled for 10 min at 100°C. Proteins were resolved by SDS-PAGE and transferred onto a nitrocellulose membrane, which was blocked with 5% milk in PBS-T. FOXA1 was detected with a polyclonal rabbit anti-FOXA1 antibody (ChIPAb+™ FOXA1 Antibody, Sigma-Aldrich, 17-10267) and, as a loading control, β-actin was detected with a mouse anti-β-actin antibody (abcam, ab8227), followed by incubation with appropriate secondary antibodies (Jackson ImmunoResearch). SuperSignal^TM^ West Pico PLUS chemiluminescent substrate (Thermo Fisher Scientific) was applied and membranes imaged with the Invitrogen^TM^ iBright^TM^ imaging system. ImageJ software was used to quantify protein levels.

### 2.4 ChIP-qPCR

Chromatin was extracted using a method adapted from Ghisletti *et al*. 2010.^[41]^ 5×10^6^ Apollo cells were fixed with 16% formaldehyde, lysed and then sonicated with the BioruptorPlus (Diagenode) using 10 cycles of [30 s “ON”, 30 s “OFF”]. Immunoprecipitation used 5 µg per ChIP of antibodies: FOXA1 (Sigma-Aldrich, 17-10267); H3K27ac (abcam, ab4729); p300/KAT3B (abcam, ab275378); SMARCA2 (abcam, ab240648); H3 (abcam, ab1791) and pre-immune serum negative control (produced in-house). ChIP-qPCR primers are shown in Table 1. ChIP experiments were repeated using cell pools from three biological repeats. Each biological repeat sample was loaded in duplicate on the QuantStudio^TM^ 7 real-time PCR cycler (Thermo Fisher Scientific) with 45 amplification cycles. Enrichment was calculated in Excel and normalised to 1% of the starting chromatin (input).

**TABLE 1.**
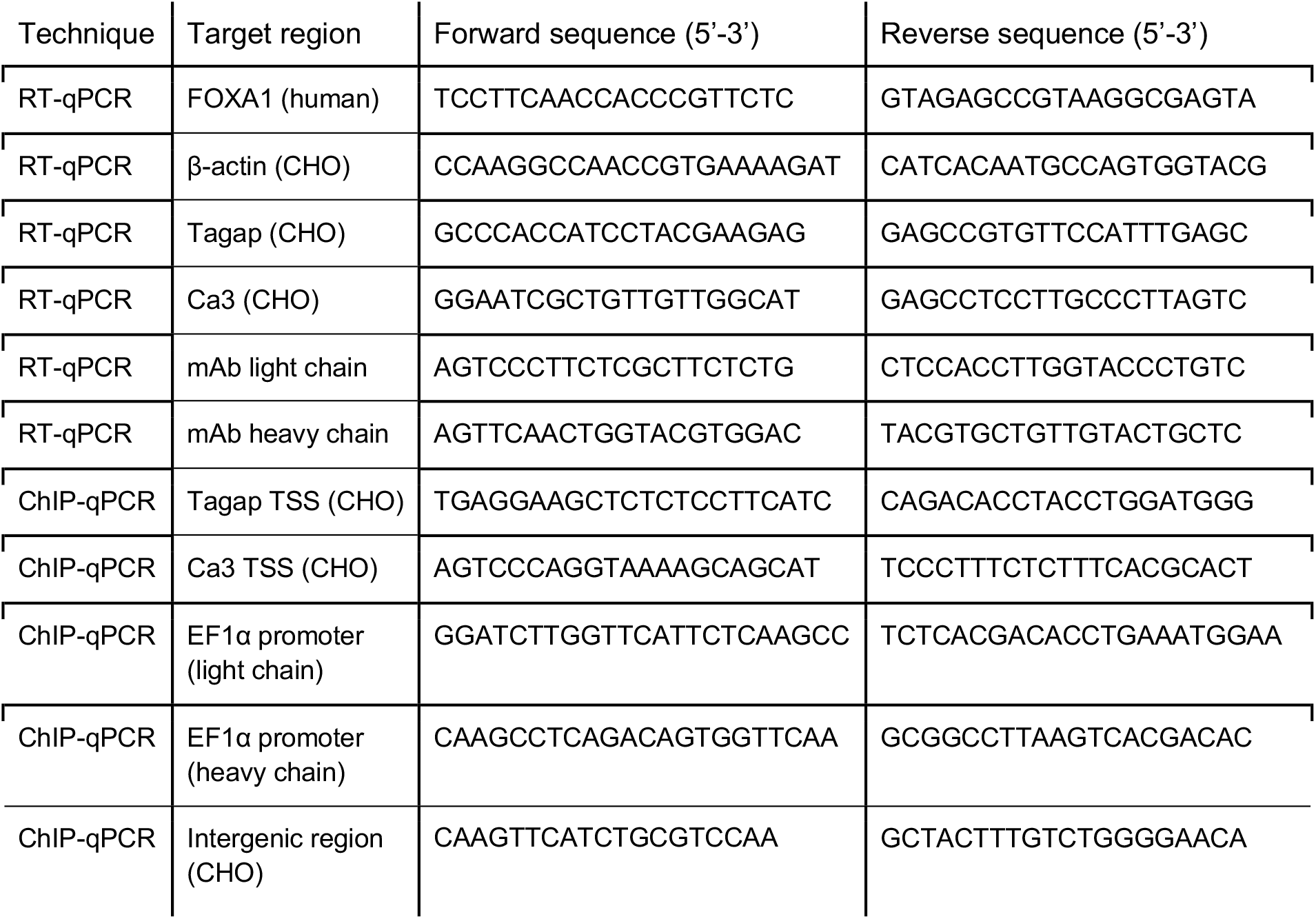
ChIP-qPCR and RT-qPCR primer sequences.

### 2.5 CpG methylation

For DNA methylation analysis, the EpiMark Methylated DNA Enrichment Kit (NEB, E2600S) was used. 5 mg genomic DNA was extracted using the Monarch^®^ Genomic DNA Purification Kit (New England BioLabs) and sonicated to 250 bp using a Covaris Focused Ultrasonicator (Covaris). Methylated DNA was isolated using Protein A magnetic beads coupled to MBD2-Fc protein. Following incubation, unbound DNA was washed off and captured CpG-methylated DNA was eluted in DNase-free water. The resulting enriched CpG-methylated DNA was quantified by qPCR with primers against target regions (Table 1). Samples were measured in duplicate on the QuantStudio^TM^ 7 real-time PCR cycler (Thermo Fisher Scientific) with 45 amplification cycles. Enrichment was calculated in Excel and values were normalised to 1% of the starting material (input).

### 2.6 mRNA expression by RT-qPCR

Total RNA from 1×10^6^ cells was isolated from stably-expressing Apollo cell pools using TRIzol reagent (Invitrogen).Concentration and quality of RNA was measured on a Nanodrop One (Thermo Fisher Scientific). cDNA was produced using the SuperScript IV Low-Input cDNA PreAmp kit (Thermo Fisher Scientific) with RNasin® Plus inhibitor (Promega) in 20 µl reactions. cDNA samples alongside RNA samples as negative controls were used as templates in a 20 µl reaction with primers against target genes (Table 1). Primer sequences for Tagap and Ca3 were obtained from Berger *et al*. 2020.^[30]^ Samples were measured in duplicate on the QuantStudio^TM^ 7 real-time PCR cycler (Thermo Fisher Scientific) with 45 amplification cycles. Relative transcript levels were normalised to β-actin.

### 2.7 ELISA

Production of mAb was quantified by ELISA. Apollo cell pools overexpressing FOXA1 or control plasmids were seeded at 0.5×10^6^ per well in 25 cm^2^ culture flasks in 15 ml media. Samples of culture medium were collected 24-hours post-seeding and cells were removed. 96-well plates coated with Protein A were washed (PBS, 0.05% Tween20). mAb-containing samples and purified human IgG were serially diluted (PBS, 0.05% Tween20), added in triplicate into the wells of the 96-well plate and incubated for 1h at RT. Plates were washed (PBS, 0.05% Tween20) and chicken pAb to human IgG HRP (abcam, ab112454) in Superblock^TM^ Blocking Buffer (Thermo Fisher Scientific) was transferred into all wells and incubated for 1h at room temperature. After washing, detection agent SuperSignal^TM^ ELISA Pico Chemiluminescent Substrate (Thermo Fisher Scientific) was added and mAb levels quantified on a CLARIOstar Plate Reader (BMG Labtech).

## 3 Results

### 3.1 FOXA1 can bind Tagap, Ca3 and EF1α promoters in CHO cells

The EF1α promoter drives high expression of the eukaryotic translation elongation factor 1 alpha 1 (EEF1A1) gene in human cells and has been used in an industrial capacity to achieve robust, constitutive expression of transgenes.^[33,34]^ Motif enrichment analysis revealed two adjacent recognition sequences for FOXA1 in the human EF1α promoter sequence (Figure 1A). Although motif analysis identifies sequences that are predicted to be bound by a transcription factor, such sites are often inaccessible *in vivo* due to chromatin folding and/or occlusion by other proteins. We therefore used chromatin immunoprecipitation (ChIP) to test if the human EF1α promoter can be bound by FOXA1 protein in CHO cells in a context relevant to industrial use. The system tested was the Apollo^TM^ platform, a DG44-derived CHO cell line optimised for industrial use. A clone was tested with stably-integrated mAb light and heavy chain transgenes driven by EF1α promoters (Figure 1B). When an expression vector encoding HA-tagged FOXA1 was transfected into these cells, levels of FOXA1 protein increased 2.9-fold (p<0.01) compared to cells transfected with empty vector (Figure 1C). ChIP-PCR demonstrated that binding of FOXA1 at the EF1α promoter increased 6-fold (p<0.01) and 3-fold (p<0.001) at the mAb light and heavy chain genes, respectively (Figure 1D).

**FIGURE 1.**
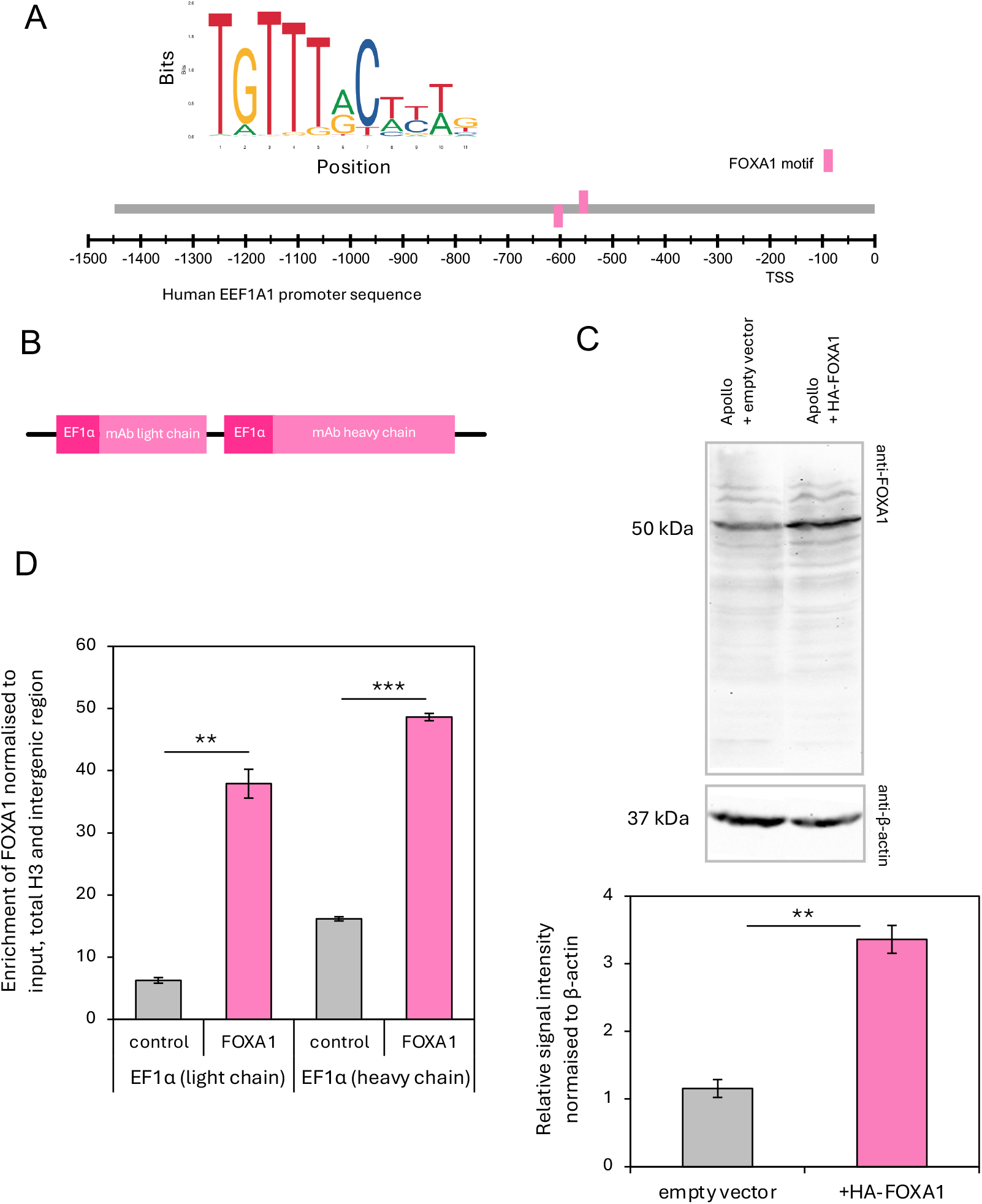
FOXA1 can bind EF1α promoters in CHO cells. (A) FOXA1 consensus binding motif (top) and motif enrichment analysis of FOXA1 binding sequences in the EF1α promoter (bottom). (B) Schematic of construct containing EF1α-driven mAb light and heavy chain genes stably expressed in CHO cells. (C) Representative western blot showing FOXA1 and β-actin levels in Apollo cells transfected with empty vector or with HA-FOXA1 (top). Quantification of FOXA1 relative to β-actin in western blots (bottom). (D) ChIP data showing enrichment of FOXA1 at the EF1α promoter in Apollo cells transfected with empty vector or HA-FOXA1. Data is normalized to input, total H3 and a negative control region. ^**^p<0.01, ^***^p<0.001 (*t*-test). Error bars represent ± SEM, *n=3*.

Overexpression of FOXA1 in CHO-M cells has been shown to induce the endogenous Tagap and Ca3 genes, which can stimulate yields of transgene products.^[30]^ Whether this reflected direct binding was not established, but our motif analysis identified multiple FOXA1 target motifs in the Tagap and Ca3 promoters (Figure 2A). We used ChIP-PCR to test if these genes are targeted directly upon FOXA1 overexpression. Indeed, FOXA1 binding was clearly enriched in both cases (p<0.05; Figure 2B). Together these data demonstrate that, when overexpressed in CHO cells, FOXA1 can act directly on promoters driving transcription of transgenes, as well as endogenous genes shown to have beneficial effects on productivity.

**FIGURE 2.**
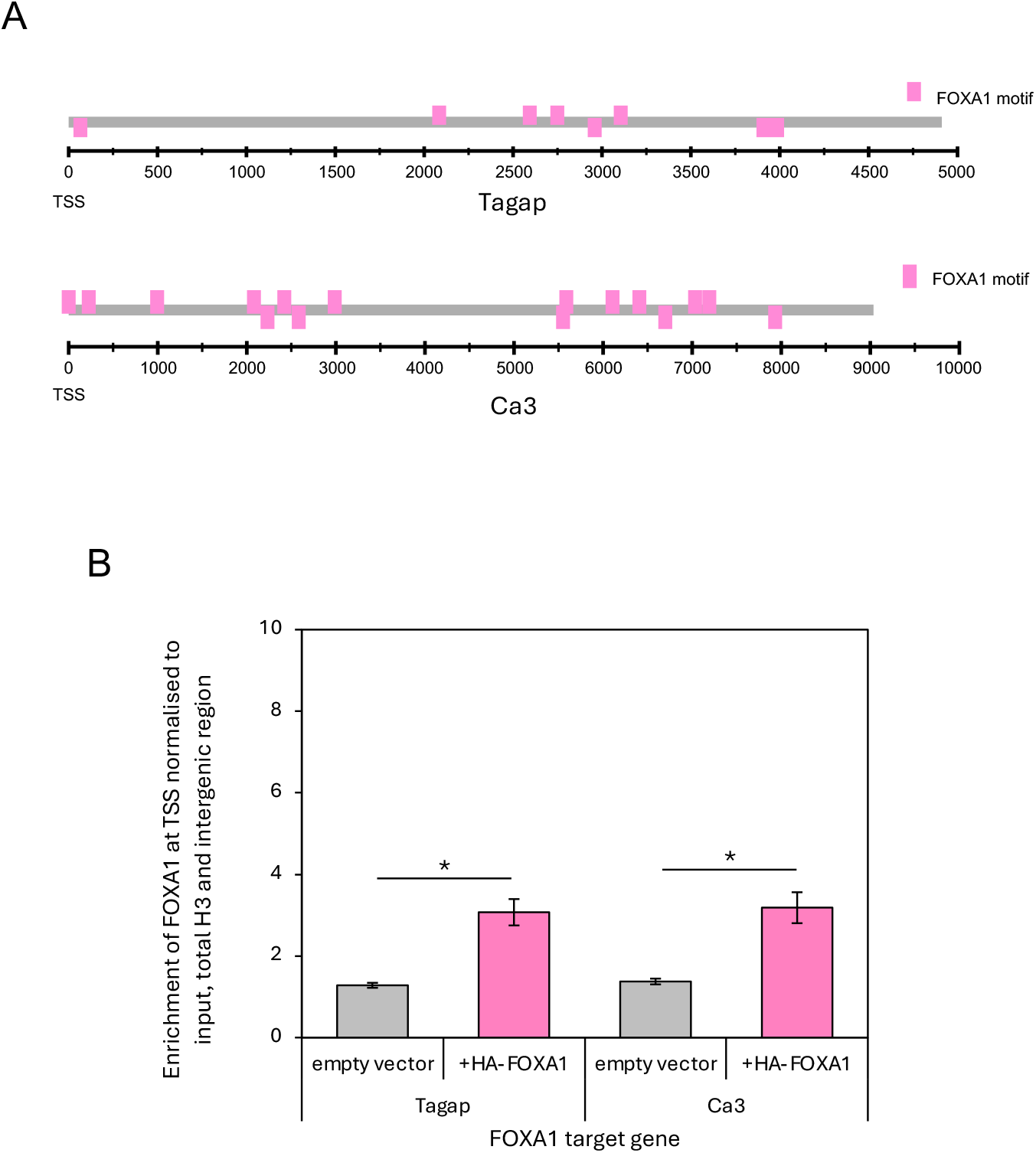
FOXA1 can bind Tagap, Ca3 in CHO cells. (A) Motif enrichment analysis of FOXA1 binding at the promoters of CHO Tagap (top) and Ca3 (bottom) genes. (B) ChIP-PCR data showing enrichment of FOXA1 at the Tagap and Ca3 promoters in Apollo cells transfected with empty vector or HA-FOXA1. Data are normalized to input, total H3 and a negative control region. ^*^p<0.05 (*t*-test). Error bars represent ± SEM, *n=3*.

### 3.2 FOXA1 overexpression can improve the epigenetic environment of target promoters

FOXA1 occupancy of regulatory elements facilitates maintenance of the epigenetic signatures of active transcription. This is achieved through recruitment of cofactors, such as the important histone acetyltransferase p300 (EP300).^[35]^ ChIP demonstrated binding of endogenous p300 to the transgenic EF1α promoters in Apollo cells (Figure 3A). FOXA1 overexpression caused further increases in p300 occupancy, of 2.7-fold at the light chain (p<0.05) and 1.4-fold at the heavy chain mAb gene (p>0.05). Acetylation of histone H3 on lysine 27 (H3K27ac) is catalysed by p300 and, accordingly, is enriched significantly at the EF1α promoters for the light and heavy chain genes (2.7-fold, p<0.01, and 1.7-fold, p<0.05, respectively), when FOXA1 is overexpressed (Figure 3B). These effects are selective, as p300 and H3K27ac did not increase at three other loci tested in parallel (Figures 3A-B).

**FIGURE 3.**
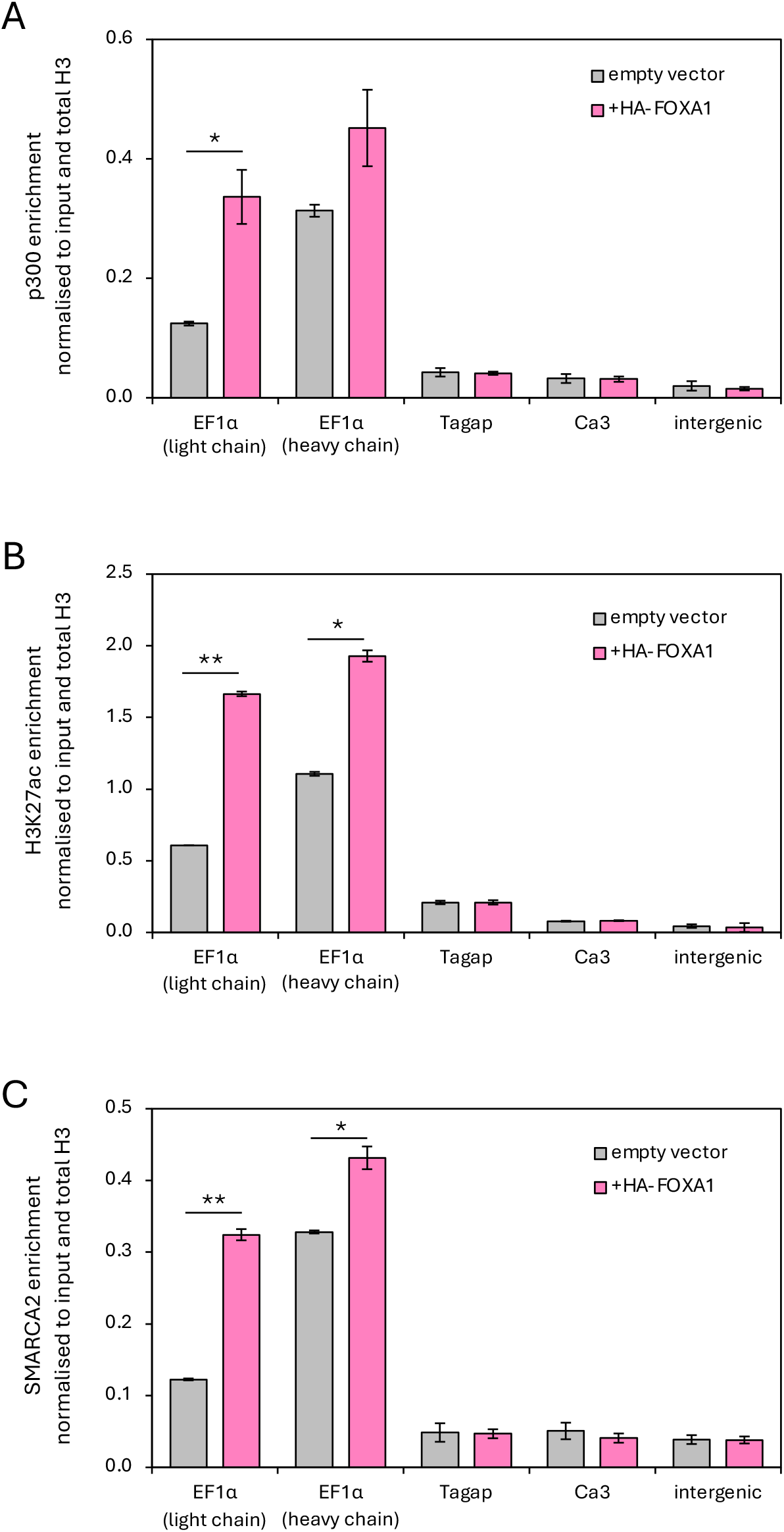
FOXA1 overexpression can improve the epigenetic environment of target promoters. (A-C) ChIP data showing enrichment of p300 (A), H3K27ac (B) and SMARCA2, (C) at EF1α, Tagap and Ca3 promoters and an intergenic region, normalised to input data and total H3. ^*^p<0.05, ^**^p<0.01 (*t*-test). Error bars represent ± SEM, *n=3*.

Promotion of transcription following chromatin opening by FOXA1 has been associated with recruitment and activity of the SWI/SNF chromatin remodelling complex, which binds to p300.^[22,35,36]^ ChIP using an antibody against SMARCA2, one of the ATPase subunits of the SWI/SNF complex, revealed significantly increased enrichment at the light and heavy chain EF1α promoters upon FOXA1 overexpression (2.6-fold, p<0.01 and 1.3-fold, p<0.05, respectively; Figure 3C). This supports a model in which recruitment of SWI/SNF may provide an additional mechanism of transcriptional activation by FOXA1 in this context.

Contrary to expectations, p300 and SMARCA2 occupancy failed to increase at the Tagap and Ca3 promoters, when tested in parallel (Figures 3A and 3C), despite the significantly elevated FOXA1 binding at these sites (Figure 2B). Consistent with the lack of p300 recruitment, the relatively low levels of H3K27ac at these promoters also remained unchanged when FOXA1 is overexpressed (Figure 3B).

STRING reveals that FOXA1 also interacts with TET1, a dioxygenase that catalyses removal of repressive 5-methylcytosine modifications from DNA (Figure 4A).^[35]^ Consistent with this, FOXA1 overexpression results in reduced levels of 5-methylcytosine at the transgenic EF1α promoters (1.7-fold, p<0.05 and 1.6-fold, p<0.01, respectively), and also at the endogenous promoters of the Tagap and Ca3 genes (2.0-fold, p<0.05 and 1.5-fold, p<0.05, respectively). This effect is specific, as it is not observed at the control intergenic region where FOXA1 fails to bind (Figure 4B). Collectively, these data show that overexpressing FOXA1 in CHO cells can selectively elicit several epigenetic changes that are conducive to transcription (Figure 4C).

**FIGURE 4.**
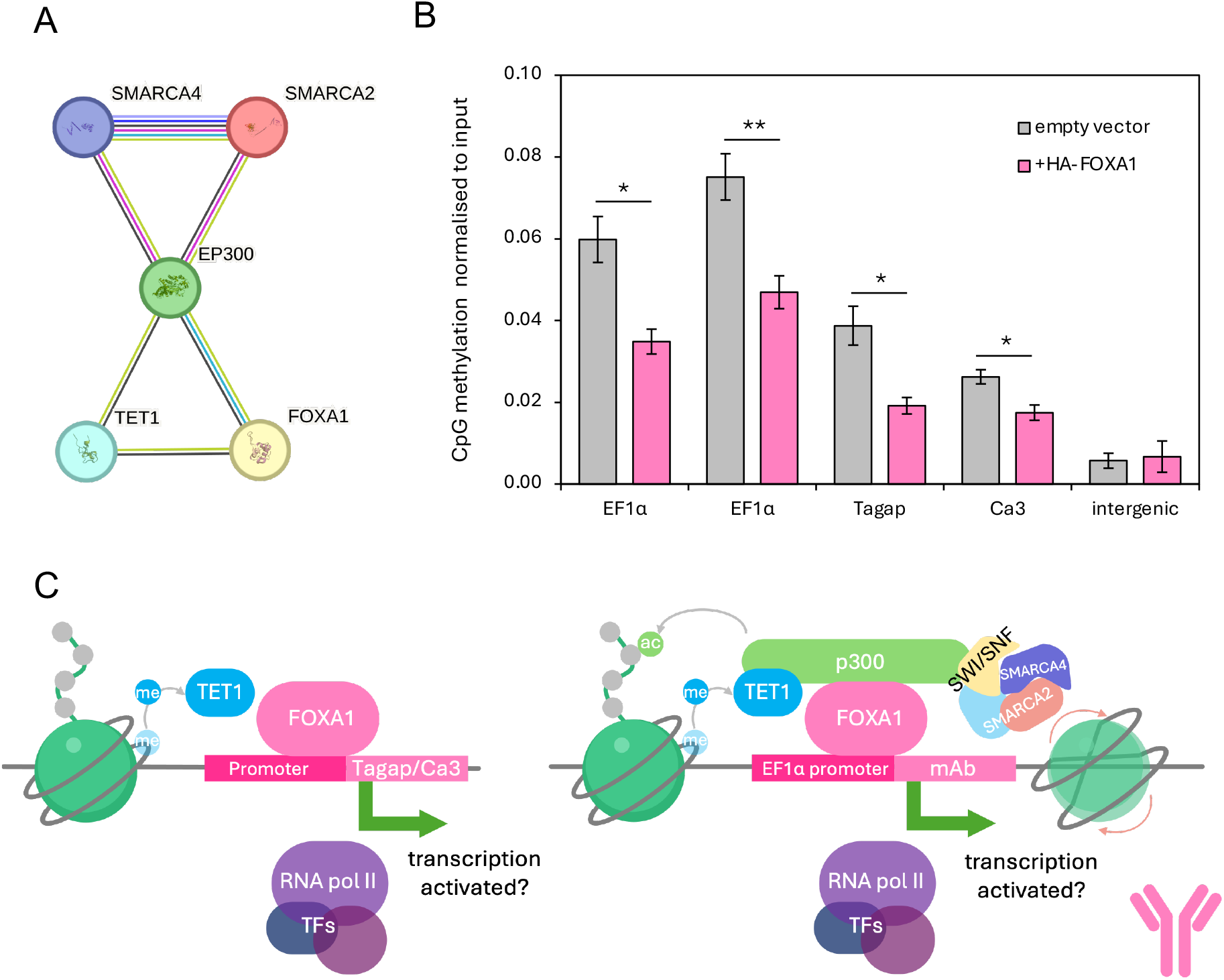
FOXA1 occupancy elicits further epigenetic changes. (A) STRING interaction network of FOXA1, TET1, p300, and the SWI/SNF proteins SMARCA2 and SMARCA4. (B) DNA (CpG) methylation at the EF1α, Tagap and Ca3 promoters and intergenic region in Apollo cells transfected with empty vector or HA-FOXA1. Data normalised to input. ^*^p<0.05, ^**^p<0.01 (*t*-test). Error bars represent ± SEM, *n=3*. (C) Schematic depicting FOXA1 binding target promoters and inducing demethylation of CpG (left) and, in some cases, recruiting additional epigenetic modifiers (right).

### 3.3 Overexpression of FOXA1 improves productivity of Apollo cells

To investigate the impact of increased FOXA1 on expression, RT-qPCR analyses were conducted. As observed by Berger *et al*. (2020) in CHO-M cells,^[30]^ significant upregulation of the endogenous Tagap and Ca3 genes was induced by FOXA1 overexpression in Apollo cells, with mRNA levels elevated by 3.0-fold (p<0.05) and 2.0-fold (p<0.01), respectively (Figure 5A). MAb light and heavy chain mRNAs were also significantly elevated under these conditions, by 2.8-fold (p<0.01) and 2.2-fold (p<0.01), respectively (Figure 5B). It is noteworthy that induction of the transgenes was no greater than that of the endogenous Tagap gene, despite the additional epigenetic regulators that were detected.

**FIGURE 5.**
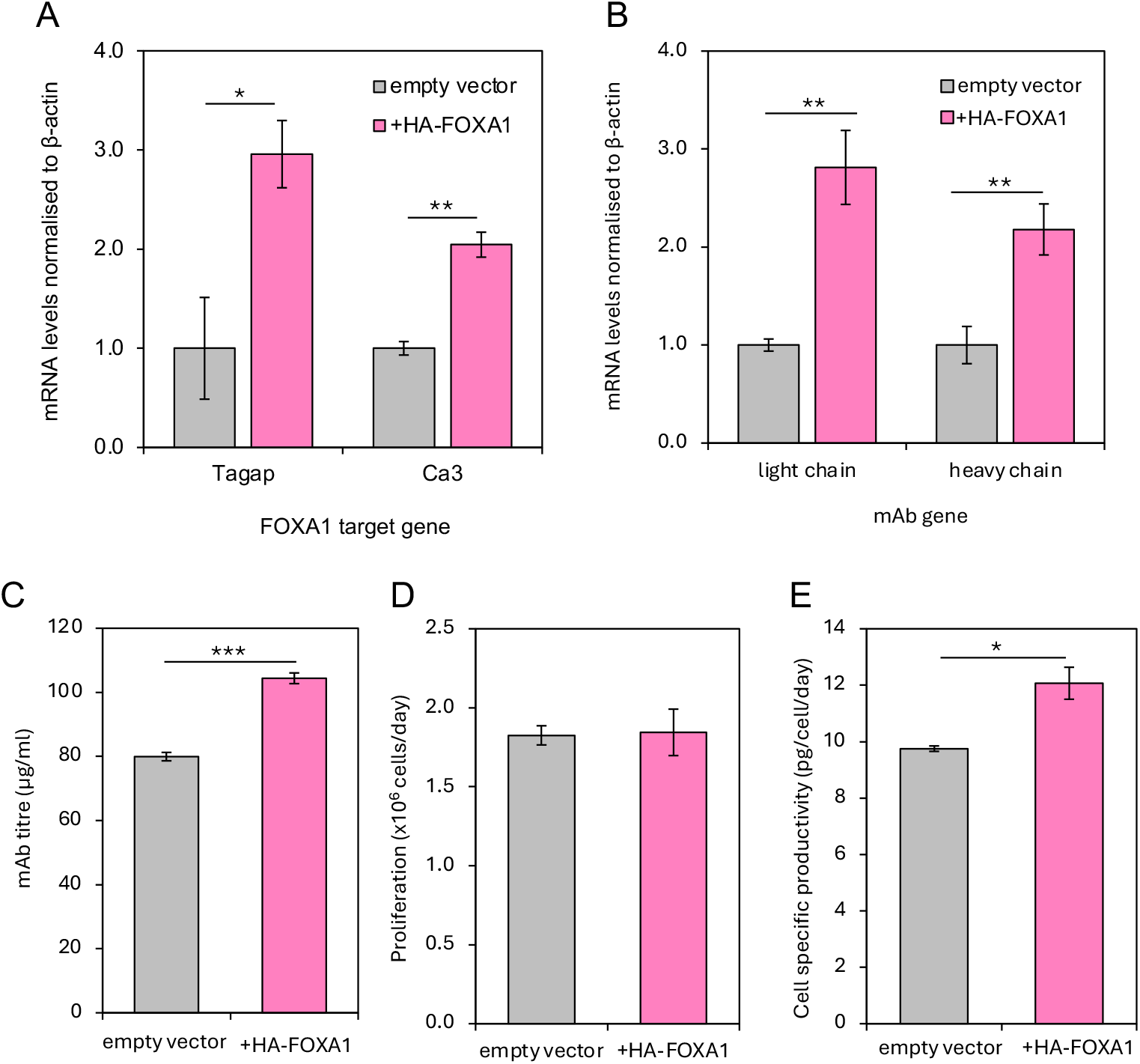
Overexpression of FOXA1 improves productivity of Apollo cells. (A) Tagap and Cas9 mRNA levels, (B) mAb light and heavy chain mRNA levels, (C) mAb titre, (D) cell proliferation and (E) cell specific productivity in Apollo cells transfected with empty vector or HA-FOXA1. ^*^p<0.05, ^**^p<0.01, ^***^p<0.001 (*t*-test). Error bars represent ± SEM, *n=3*.

We then measured the titre of secreted antibody and found it to be elevated significantly (1.2-fold, p<0.001, Figure 5C). This increase was not due to altered rates of cell proliferation (Figure 5D). A 20% higher specific productivity was obtained (Figure 5E), providing an additional 2.4 pg/cell/day (p<0.01). Overexpression of FOXA1 can therefore raise mAb mRNA, titre and specific productivity in a CHO cell line that is optimised for industrial use.

## 4 Discussion

These findings illustrate the multifaceted impact of the pioneer factor FOXA1 in determining chromatin organisation and transcriptional activity. They also demonstrate its potential for use in biotechnology as a means to enhance productivity. Direct binding of FOXA1 at the EF1α promoter can substantially increase transcription of mAb transgenes via a cascade of complementary epigenetic changes. These effects are supported by the selective induction of endogenous genes which improve the cells’ capacity for production. As a consequence, overexpression of FOXA1 can selectively elevate the specific productivity of the optimised Apollo CHO cell line.

Our data suggest that the epigenetic state of the human EF1α promoter linked to transgenes is suboptimal in Apollo cells for maximal transcription. This is noteworthy, as the EF1α promoter is widely used and considered highly efficient. Nevertheless, we find that its efficacy can be raised using supraphysiological FOXA1 levels. The important coactivator p300 can be recruited to this promoter by FOXA1 to cause local enrichment of H3K27ac, an established marker of active promoters. In addition to acetylating H3K27, p300 also interacts with the SWI/SNF remodelling complex, that can displace nucleosomes to enhance DNA accessibility. Promoter utilisation can also be increased by the DNA demethylase TET1, which interacts with FOXA1 and removes inhibitory methyl groups to facilitate transcription.^[11,24]^

We previously used synthetic CRISPR-dCas9 systems to establish that p300 and TET1 can act directly to raise the titre of mAb secreted from Apollo cells.^[11,24]^ This was achieved by targeting dCas9-p300 and dCas9-TET1 with specific guide RNAs to CMV promoters driving mAb transgenes. Here we have shown that the same effects can be achieved using FOXA1, which offers a more practical approach than the combined use of two large dCas9 fusion proteins.

It is notable that increased binding of FOXA1 does not elicit the same epigenetic changes at two other promoters. Although FOXA1 enrichment at the promoters of Tagap and Ca3 raises expression of these genes, it is not accompanied by concurrent increases in local p300, H3K27ac or SMARCA2. This suggests that a FOXA1-induced reduction in CpG methylation may be sufficient for transcriptional activation in these contexts. Alternatively, induction of these genes by FOXA1 may involve recruitment of other cofactors that were not assayed in this study. It is unclear why p300 and SMARCA2 are recruited to exogenous EF1α promoters but not to the promoters of endogenous Tagap and Ca3 genes. Neither p300 nor SWI/SNF complexes containing SMARCA2 recognise specific DNA sequence motifs that might confer promoter selectivity; instead, they are recruited by protein-protein interactions with DNA-binding factors, in this case FOXA1. The data suggest that context influences recruitment or retention of cofactors by FOXA1 in ways that have yet to be fully characterised. In support of this, the light and heavy chain mAb transgenes show quantitative differences in their recruitment of FOXA1 and its associated factors and in their transcriptional responses, despite being driven by identical EF1α promoters. This variation is likely to reflect differences in the DNA sequences upstream and downstream of the promoters themselves, which impact chromatin context and hence gene transcription.

Although the induction of mRNA obtained was more than 2-fold, the 1.2-fold increase in cell-specific productivity was less substantial. This diminution of response may be caused by downstream bottlenecks in translation, assembly, processing or secretion of mAb.^[37]^ Nevertheless, the 20% boost in titre is significant and valuable.

Our findings demonstrate that increased titre and specific productivity of recombinant Apollo cells can be achieved by introducing exogenous FOXA1, which raises expression of transgene mRNA. Overexpression of FOXA1 was shown previously to increase production of trastuzumab and infliximab in CHO-M cells, an effect that was attributed to upregulation of the FOXA1 target genes Tagap and Ca3, as the transgenes themselves were not responsive.^[30]^ Furthermore, the authors found that overexpressing Tagap alone, but not Ca3, was sufficient to recapitulate the increase in trastuzumab achieved by transfecting FOXA1.^[30]^ In contrast, Tagap overexpression in the same study failed to stimulate as efficiently as FOXA1 the titre of infliximab.^[30]^ Thus, whilst the upregulation of endogenous Tagap and Ca3 genes, an effect also observed in our system, can contribute to increased titre through beneficial effects on the host cells, additional effects of FOXA1 can improve productivity. We have shown that FOXA1 binds to exogenous EF1α promoters driving transgenes and induces epigenetic changes that upregulate mAb mRNA production, thereby raising output more directly. Concomitant stimulation of Tagap and Ca3 can optimise metabolism and stress resistance to improve cell capacity for superior production.

## 5 Conclusion

Our findings, using an industry-standard CHO platform, support previous work in an orthogonal CHO system demonstrating beneficial effects on titre and productivity of overexpressing FOXA1. We have shown that the EF1α promoter, that is widely used to drive transgene transcription, is bound directly by FOXA1 in cells, leading to recruitment of epigenetic modifiers that remove inhibitory DNA methylation and stimulate activating histone acetylation. By employing complementary molecular mechanisms at promoters of transgenes and specific endogenous genes, FOXA1 has potential to enhance industrial production of biopharmaceutical proteins.

## Author Contributions

**Sienna P. Butterfield:** conceptualisation, methodology, verification, formal analysis, investigation, writing – original draft, visualisation | **Fay L. Saunders:** supervision, writing–review and editing | **Robert J. White:** conceptualization, writing–review and editing, supervision, project administration, funding acquisition.

## Funding

Sienna Butterfield was supported by a studentship from the Centre of Excellence in Bioprocessing 2.0 of FUJIFILM Biotechnologies UK.

## Notes

### Competing Interest Statement

The authors have declared no competing interest.

